# Identification of ADAMTS19 as a novel retinal factor involved in ocular growth regulation

**DOI:** 10.1101/2020.06.03.132852

**Authors:** Swanand Koli, Cassandre Labelle-Dumais, Yin Zhao, Seyyedhassan Paylakhi, K Saidas Nair

## Abstract

Refractive errors are the most common ocular disorders and are a leading cause of visual impairment worldwide. Although ocular axial length is well established to be a major determinant of refractive errors, the molecular and cellular processes regulating ocular axial growth are poorly understood. Mutations in genes encoding the PRSS56 and MFRP are a major cause of nanophthalmos. Accordingly, mouse models with mutations in the genes encoding the retinal factor PRSS56 or MFRP, a gene predominantly localized in the retinal pigment epithelial (RPE) exhibit ocular axial length reduction and extreme hyperopia. However, the precise mechanisms underlying PRSS56- and MFRP-mediated ocular axial growth remain elusive. Here, we show that *Adamts19* expression is significantly upregulated in retina of mice lacking either *Prss56* or *Mfrp*. Using a combination of genetic approaches and mouse models, we show that while ADAMTS19 is not required for ocular growth during normal development, its inactivation exacerbates ocular axial length reduction in both *Prss56* or *Mfrp* mutant mice. These results suggest that the upregulation of retinal *Adamts19* expression is part of an adaptive molecular response to counteract impaired ocular growth. Using a complementary genetic approach. We further demonstrate that loss of PRSS56 or MFRP function prevents excessive ocular axial growth in a mouse model of developmental myopia caused by a null mutation in *Irpb*, demonstrating that ocular axial elongation in *Irbp*^*-/-*^ mice is fully dependent on PRSS56 and MFRP functions. Collectively, our findings provide insight into the molecular network involved in ocular axial growth regulation and refractive development and support the notion that relay of the signal between the retina and RPE could be critical for promoting ocular axial elongation.

## INTRODUCTION

Nanopthalmos is a rare developmental disorder characterized by significantly smaller but structurally normal eyes and extreme hyperopia resulting from compromised ocular growth [1]. Also, nanophthalmic individuals are highly susceptible to developing blinding conditions including secondary angle-closure glaucoma, spontaneous choroidal effusions, cataracts, and retinal detachment [1]. Both sporadic and familial forms of napophthalmos with autosomal dominant or recessive inheritance have been reported [2]. To date, six genes (*PRSS56, MFRP, TMEM98, CRB1, BEST1*, and *MYRF*) have been implicated in familial forms of nanophthalmos, with PRSS56 and MFRP mutations accounting for the most frequent causes among multiple cohorts [1, 3-10]. Furthermore, the eyes of nanophthalmic individuals with biallelic mutations in *PRSS56* or *MFRP* were found to be significantly smaller compared to those carrying dominant mutations in *TMEM98* or *MYRF*. Interestingly, common variants of *PRSS56* and *MFRP* have also been found to be associated with myopia, a condition phenotypically opposite to nanophthalmos that is characterized by increased ocular elongation [11]. Together, these findings underscore the importance of PRSS56 and MFRP in ocular size regulation[2].

Ocular growth can be broadly divided into two distinct phases that take place pre- and postnatally[12]. Prenatal ocular growth occurs in the absence of visual stimulation and is primarily dictated by genetic factors [13]. In contrast, postnatal ocular growth also referred to as emmetropization, is a vision-guided process modulated by the refractive status of the eye to ensure that the axial length matches the optical power of the eye to achieve optimal focus and clear vision. Abnormal postnatal ocular axial growth leading the increased axial length constitutes a major cause of myopia, a condition characterized by blurred vision caused by focused images falling in front of the retina [12, 14, 15]. Nanophthalmos is generally attributed to impaired prenatal ocular growth as individuals with this condition are born hyperopic [1, 16]. Interestingly, in addition to being responsible for nanophthalmos, common variants of *MFRP* and *PRSS56* have also been found to be associated with myopia in the general population (an opposite condition) that primarily results from alterations in postnatal ocular axial growth [11, 17]. Thus, the association of PRSS56 and MFRP with nanophthalmos and myopia support a role for these factors in the regulation of embryonic and postnatal ocular growth development and suggest that the molecular mechanisms underlying pre- and postnatal ocular growth are shared.

It is generally accepted that postnatal ocular growth is regulated by a cascade of signaling events by which information is relayed from the retina to the sclera to induce scleral extracellular matrix (ECM) remodeling to promote ocular axial elongation and [14, 18]. Notably, PRSS56 expression is specifically detected in the retina [13], which is consistent with a central role for the retina in ocular growth regulation. MFRP is predominantly expressed in the retinal pigment epithelium (RPE) and ciliary epithelium[16] and is implicated in the transmission of molecular cues between retina and sclera during ocular growth. Using a genetic mouse model, we have recently demonstrated that the genetic ablation of *Prss56* from retinal Müller glia leads to a significant reduction in ocular axial length and hyperopia[13]. Similarly, mice and zebrafish lacking MFRP exhibit ocular axial length reduction, and *MFRP* variants in humans are associated with myopia[19-21].

Although current shreds of evidence support a key role of MFRP and PRSS56 in ocular axial length determination, the underlying mechanisms remain elusive. In this study, we use *Prss56* and *Mfrp* mutant mouse model in combination with complementary genetic approaches to gain insights into the molecular network involved in ocular size regulation. Importantly, we identified characteristic changes in retinal gene expression in response to impaired ocular growth. Specifically, we show that *Adamts19* mRNA levels are significantly increased in the retina of *Prss56* and *Mfrp* mutant mice and provide evidence that the upregulation of retinal *Adamts19* expression is part of an adaptive molecular response to impaired ocular growth. Furthermore, we demonstrate that loss of PRSS56 or MFRP function prevents excessive ocular axial elongation in a mouse model of early-onset myopia caused by a mutation in *Irbp*. Collectively, our finding hints at a potential molecular link between Müller glia and RPE involved in ocular axial growth regulation.

## RESULTS

### *Adamts19* expression is upregulated in the retina of *Prss56* mutant mice

To begin addressing the molecular processes underlying PRSS56-mediated ocular size regulation, we performed RNA-Seq analysis on the retina from *Prss56*^*gclr4*^ mutant mice and their wild-type littermates. We recently demonstrated that the ocular size reduction that we originally described in mice with a *Prss56*^*gclr4*^ mutation (causing PRSS56 protein truncation) result from a loss of function mechanism, hence, *Prss56*^*gclr4/gclr4*^ mice will be referred to as *Prss56*^*-/-*^ throughout the manuscript for simplicity[13]. Our transcriptome analysis identified *Prss56* and *Adamts19* as the top two differentially expressed genes between *Prss56* mutant (*Prss56*^-/-^) and control (*Prss56*^+/-^) retina. Consistent with the RNA-Seq data, the qPCR analysis revealed that *Prss56* and *Adamts19* mRNA levels were significantly upregulated in the retina of *Prss56* mutant mice (*Prss56*^*-/-*^) compared to their *Prss56*+^*/-*^ *and Prss56*^+*/*+^ littermates at both ages examined (postnatal day (P) 15 and P30) (Fig. 1A-B). Importantly, *Prss56* and *Adamts19* retinal expression levels in heterozygous *Prss56*^+*/-*^ mice were comparable to those detected in *Prss56*^+*/*+^ mice, which is consistent with the absence of an ocular phenotype in *Prss56*^+*/-*^ mice [13]. *Prss56*^+*/-*^ mice were therefore used as controls for all experiments presented in this study. As described previously [13], we detected a progressive upregulation of *Prss56* mRNA levels in *Prss56*^*-/-*^ retina from P15 to P60 (Fig. S1). The increase in *Adamts19* retinal expression of was found to precede that of *Prss56* in *Prss56*^*-/-*^ retina, and was detected as early as P10 and gradually increased to reach peak expression levels by P30 (Fig. S1). Notably, qPCR-Ct values suggested that the expression of *Adamts19* was minimal or negligible in *Prss56*^+*/*+^ and *Prss56*^+*/-*^ retina. Furthermore, the upregulation of retinal *Prss56* and *Adamts19* expression was also observed in mice carrying a null allele of *Prss56* (*Prss56*^*Cre*^), which we had described previously [13]. Thus, confirming that the increase in *Prss56* and *Adamts19* expression results from a loss of PRSS56 function (Fig. 1C). To determine the spatial distribution of *Adamts19* mRNA, we next performed *in situ* hybridization on ocular sections from *Prss56*^*-/-*^ and *Prss56*^+*/-*^ mice. Despite using the highly sensitive QuantiGene View RNA *in situ* hybridization method, *Adamts19* expression was only detected in *Prss56*^*-/-*^ retina, indicating that *Adamts19* expression was below the threshold of detection in control *Prss56*^+*/-*^ retina (Fig. 1D). In *Prss56* mutant retina, *Adamts19* expression was predominantly observed in the inner nuclear layer (INL), a region containing the cell bodies of Müller glia, a cell type in which *Prss56* is normally expressed. Collectively, these findings demonstrate that in addition to causing ocular size reduction, loss of PRSS56 function leads to alterations in retinal gene expression marked by increased *Prss56* and *Adamst19* mRNA levels.

**Figure 1.**
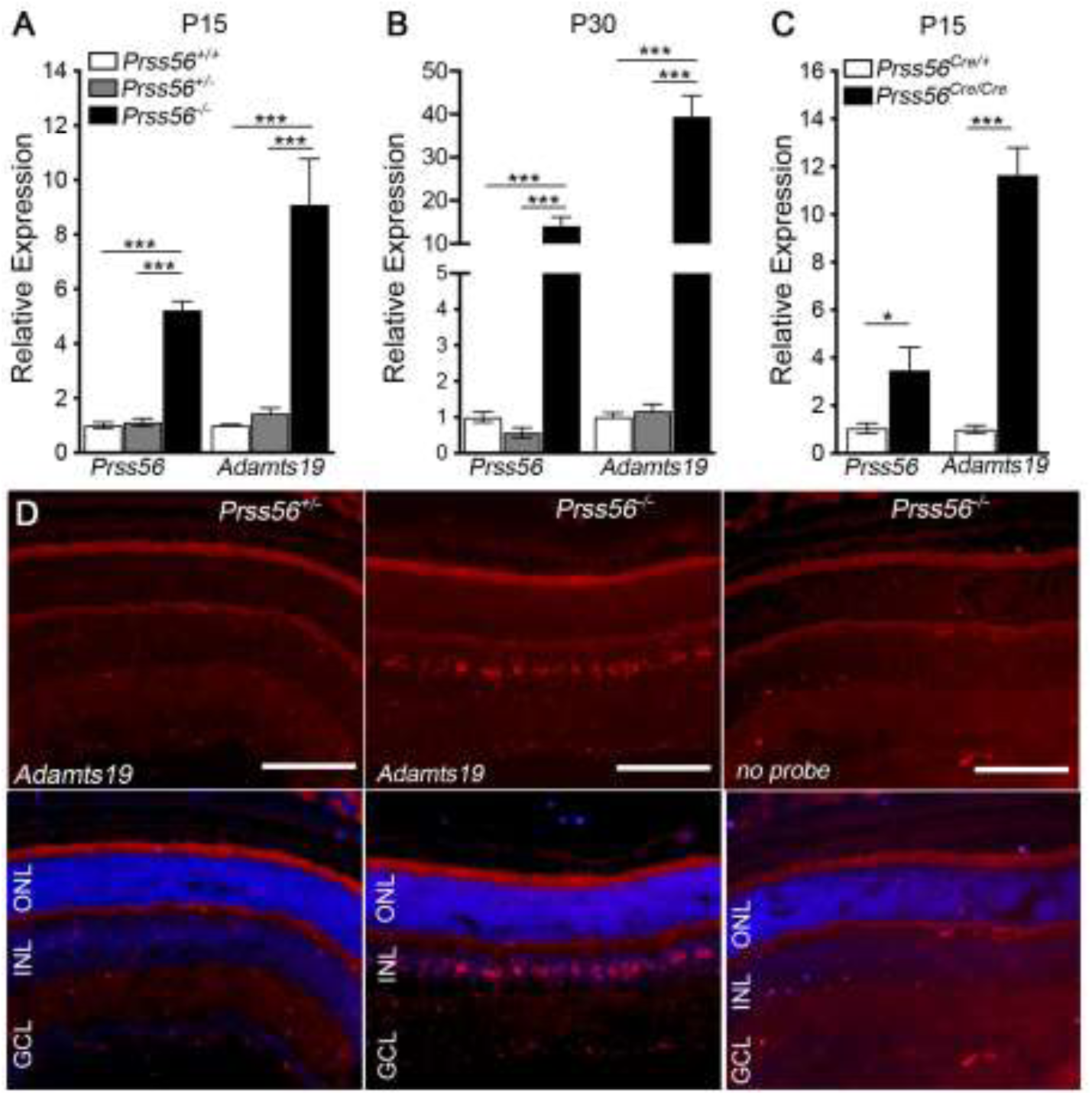
*Adamts19* expression is upregulated in the retina of *Prss56* mutant mice. (**A**-**C**) Graph showing quantification of *Prss56* and *Adamts19* mRNA levels using qPCR in P15 (**A** and **C**) and P30 (**B**) retina from *Prss56*^*glcr4*^ (**A** and **B**) and *Prss56*^*Cre*^ (**C**) mutant strains. While no difference was observed between *Prss56*^+*/*+^ and *Prss56*^*glcr4/*+^ (*Prss56*^+*/-*^) retina, a significant increase in *Prss56* and *Adamts19* mRNA levels was detected in *Prss56*^*glcr4/glcr4*^ (*Prss56*^*-/-*^) retina compared to *Prss56*^+*/*+^ and *Prss56*^*glcr4/*+^ retina at both ages examined (**A** and **B**). Similarly, significant increases in *Prss56* and *Adamts19* mRNA levels were detected in *Prss56*^*Cre/Cre*^ retina compared to the control *Prss56*^*Cre/*+^ retina. *Prss56* and *Adamts19* expression were normalized to the expression of three housekeeping genes (*Hprt1, Actb1*, and *Mapk1)*. Data are presented as fold expression relative to wild-type (mean ± SEM), N= 4 to 6 retina/group, data are presented as mean+/-SEM, **\***p<0.05; **\*\*\***p<0.001, t-test. (**D**) QuantiGene View RNA *in situ* hybridization revealed that *Adamts19* expression was below the threshold level of detection in control *Prss56*^*gclr4/*+^ retina and was only detectable in *Prss56*^*gclr4/gclr4*^ retina at P16. *Adamts19* expression (red) was predominantly observed in the INL of *Prss56* mutant retina with low levels also detected in the GCL. Scale bars: 100μm

### Retinal Prss56 and Adamts19 mRNA levels are upregulated in response to ocular size reduction in Prss56 mutant mice

To determine whether the upregulation in retinal *Adamts19* and *Prss56* mRNA levels correlates with ocular size reduction in *Prss56* mutant mice, we took advantage of the *Egr1; Prss56* double mutant mouse model (*Egr1*^*-/-*^; *Prss56*^*-/-*^) that we described previously [13]. EGR1 (early growth response1) is a major regulator of ocular growth and *Egr1*^*-/-*^ mice exhibit increased ocular axial length[13, 22]. We have previously shown that *Egr1* inactivation rescues the reduction in ocular axial length and vitreous chamber depth (VCD) in *Prss56* mutant mice as the ocular size of *Egr1*^*-/-*^;*Prss56*^*-/-*^ mice is comparable to that of control *Egr1*^+*/-*^;*Prss56*^+*/-*^ mice [12]. Using qPCR analysis, we show that in addition to rescuing ocular axial elongation, *Egr1* inactivation also prevented the increase in retinal expression of *Prss56 and Adamts19* in *Prss56* mutant mice (compare *Egr1*^*-/-*^; *Prss56*^*-/-*^ to *Egr1*^+*/-*^; *Prss56*^*-/-*^ in Fig. 2). These findings suggest that the upregulation of retinal *Prss56* and *Adamts19* does not result from loss of PRSS56 function *per se*, but rather from its effect on ocular size.

**Figure 2.**
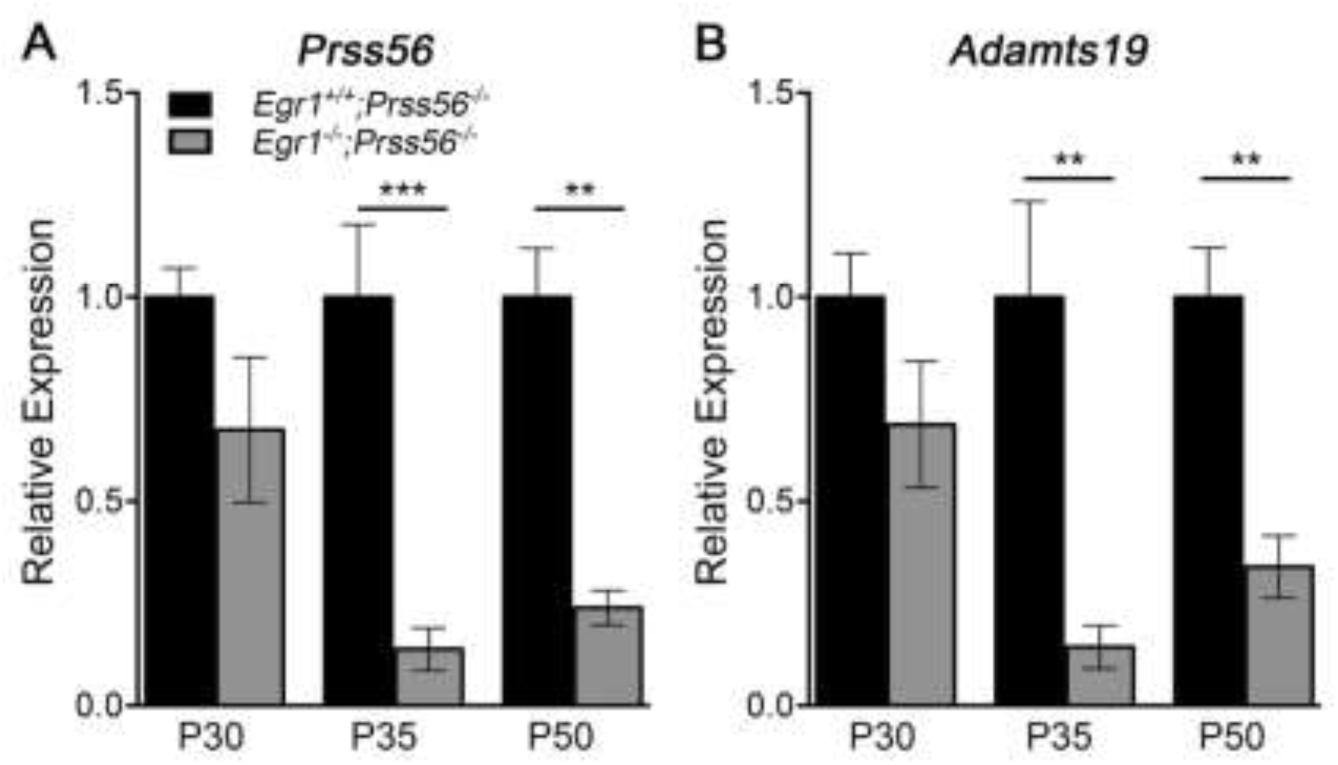
*Egr1* inactivation prevents the upregulation of retinal *Prss56* and *Adamts19* expression in *Prss56* mutant mice. (**A**-**B**) Graphs showing quantification of *Prss56* (**A**), *Adamts19* (**B**) mRNA levels using qPCR in *Prss56* mutant (*Egr1*^+*/*+^;*Prss56*^*-/-*^) and *Prss56;Egr1* double mutant (*Egr1*^*-/-*^;*Prss56*^*-/-*^) retina at different developmental stages. *Egr1* inactivation reduced retinal *Prss56* (A) and *Adamts19* (B) mRNA levels in *Prss56* mutant mice (compare *Egr1*^+*/*+^;*Prss56*^*- /-*^ *to Egr1*^*-/-*^;*Prss56*^*-/-*^). Data are presented as fold expression relative to wild-type (mean ± SEM), N=4 to 6/group. **\*\***p<0.01; **\*\*\***p<0.001, t-test.

### *ADAMTS19* is not required for ocular growth during normal development

PRSS56 and ADAMTS19 are both secreted serine proteases, raising the possibility that they might have overlapping functions in ocular growth regulation. To test this possibility, we first generated *Adamts19* knockout mice by crossing a conditional *Adamts19* mutant mouse line with the ubiquitous *β-actin-Cre line* (Fig. S2). To determine if the loss of ADAMTS19 function lead to ocular defects, we performed optical coherence tomography (OCT) to assess various ocular biometric parameters. We found that all the ocular parameters examined, including ocular axial length, VCD, and retinal thickness were indistinguishable between *Adamts19*^+*/*+^, *Adamts19*^+*/-*^ and *Adamts19*^*-/-*^ mice (Fig. 3 and Fig. S3). These findings demonstrate that ADAMTS19 is not required for ocular growth during normal development.

**Figure 3.**
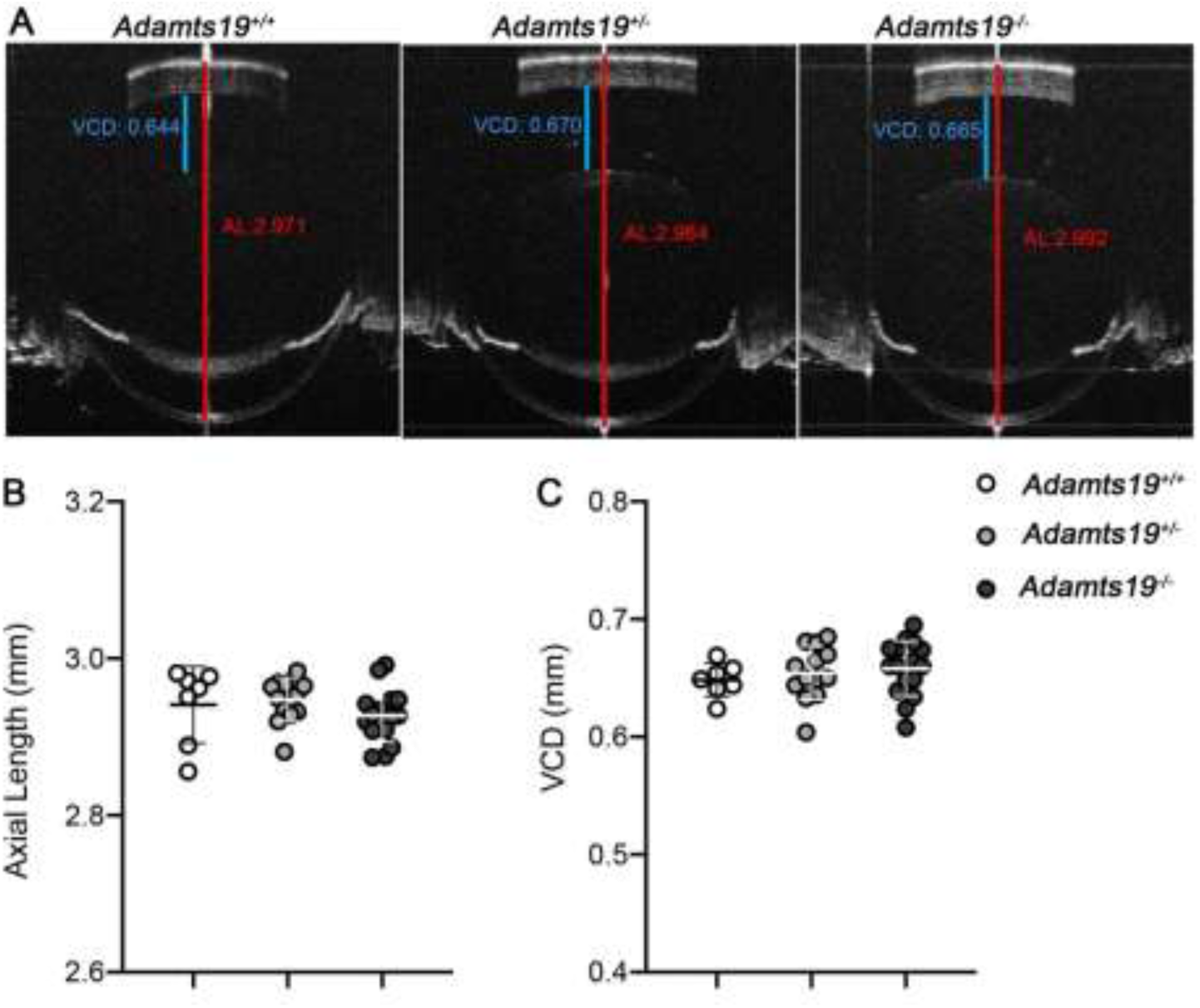
ADAMTS19 is not required for axial growth during normal ocular development. (**A**) Representative OCT images showing that ocular axial length (quantified in **B**) and VCD (quantified in **C**) are indistinguishable between *Adamts19*^*-/-*^, *Adamts19*^+*/-*^ and control *Adamts19*^+*/*+^ mice at P18. Data are presented as mean ± SD, N>7/group.

### Loss of ADAMTS19 function exacerbates ocular axial length reduction in *Prss56*^*-/-*^ mice

In light of our findings, we hypothesized that the upregulation of retinal *Adamts19* expression might be part of an adaptive molecular response to compensate for the loss of PRSS56 function and promote ocular axial growth. To this end, we tested the effect of *Adamts19* inactivation in *Prss56*^*-/-*^ mice by crossing *Prss56* mutant mice to the *Adamts19* mutant line to generate *Prss56*^*-/-*^ mice that are wild-type, heterozygous or homozygous for the *Adamts19* null allele (*Adamts19*^+*/*+^, *Adam19*^+*/-*^ or *Adamts19*^*-/-*^). Notably, since all ocular biometric parameters of *Adamts19*^+/-^; *Prss56*^+/-^ mice were comparable to those of wild-type *(Adamts19*^+*/*+^; *Prss56*^+*/*+^) littermates (Fig. S4A), *Adamts19*^+*/-*^;*Prss56*^+*/-*^ mice were used as controls. As expected, axial length and VCD were significantly reduced in all three groups of mice lacking *Prss56* (*Prss56*^*-/-*^) compared to the control mice (Fig. 4). However, the axial length and VCD were significantly reduced in *Adamts19;Prss56* double mutant mice (*Adamts19*^-/-^;*Prss56*^-/-^) compared to *Prss56* single mutants (*Adamts19*^+*/-*^; *Prss56*^*-/-*^ or *Adamts19*^+*/*+^;*Prss56*^*-/-*^) at both age examined (P18 and P30) (Fig. 4). As reported previously [13], ocular axial length reduction in *Prss56*^*-/-*^ mice was associated with an increase in retinal thickness (Fig. S5). Notably, a modest but significant increase in retinal thickness was observed in *Adamts19*^-/-^;*Prss56*^-/-^ mice compared to *Prss56* mutant mice (*Adamts19*^+*/-*^;*Prss56*^*-/-*^ and *Adamts19*^+*/*+^; *Prss56*^*-/-*^) at P18 (Fig. S5).

**Figure 4.**
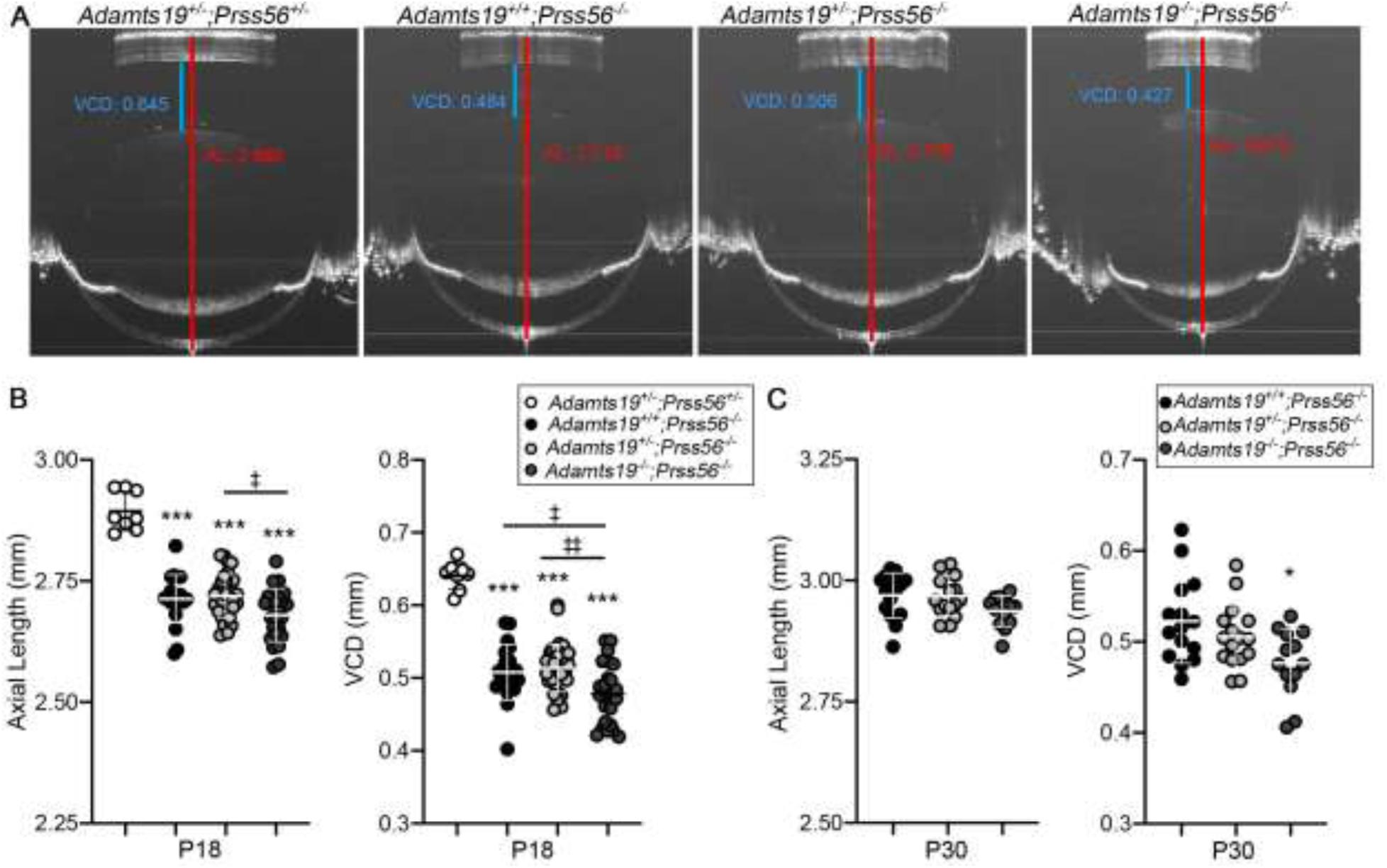
*Adamts19* inactivation exacerbates the ocular axial length reduction in *Prss56* mutant mice. (**A**) Representative OCT images showing reduced ocular axial length and VCD (quantified in **B** and **C**) in mice carrying a *Prss56* mutation (*Adamts19*^+*/*+^;*Prss56*^*-/-*^;*Adamts19*^+*/-*^;*Prss56*^*-/-*^ and *Adamts19*^*-/-*^; *Prss56*^*-/-*^) compared to control *Adamts19*^+*/-*^;*Prss56*^+*/-*^mice. Importantly, the *Adamts19;Prss56* double mutant (*Adamts19*^*-/-*^;*Prss56*^*-/-*^) mice show a modest but consistent reduction in ocular axial length and VCD compared to *Prss56* single mutant (*Adamts19*^+*/-*^;*Prss56*^*-/-*^ *or Adamts19*^+*/*+^;*Prss56*^*-/-*^) mice at both ages examined (P18 and P30). Data are presented as mean ± SD, N>13/group. **\***p<0.05; **\*\***p<0.01, One-way ANOVA.

Besides, we found that *Adamts19* expression was significantly increased in the retina from both *Adamts19*^+*/*+^;*Prss56*^*-/-*^ and *Adamts19*^+*/-*^;*Prss56*^*-/-*^ mice compared to that of *Adamts19*^+/-^;*Prss56*^+/-^ control mice, which is consistent with the observation that exacerbation of the ocular axial length reduction in *Prss56* mutant mice is only observed when *Adamts19* is completely knocked out (Fig. S5C). Together, these results demonstrate that *Adamts19* inactivation exacerbates ocular size reduction in *Prss56*^*-/-*^ mice and is consistent with the upregulation of retinal *Adamts19* expression being part of an adaptive molecular response triggered by impaired ocular growth in *Prss56* mutant mice.

### *Adamts19* inactivation exacerbates ocular axial length reduction in *Mfrp* mutant mice

Interestingly, elevated retinal levels of *Prss56* expression has recently been reported in another mouse model of nanophthalmos caused by a mutation in the gene coding for membrane frizzed related-protein (*Mfrp*) [23]. Increased *Adamts19* expression was also observed in *Mfrp*^*-/-*^ eyes but the specific ocular tissue/cell type in which *Adamts19* was expressed was not addressed [23]. Since *Adamts19* expression was specifically detected in the retina of the *Prss56*^*-/-*^ mice, we performed a qPCR analysis to confirm that the levels of *Prss56* and *Adamts19* were upregulated in the retina of *Mfrp*^*-/-*^ mice compared to control *Mfrp*^+*/-*^ littermates (Fig. 5A). To determine if *Adamts19* inactivation also exacerbates the ocular size reduction caused by *Mfrp* deficiency, we crossed *Mfrp* mutant mice with the *Adamts19* mutant line and conducted OCT analyses on the progeny. Since the ocular biometric parameters of *Adamts19*^+*/-*^; *Mfrp*^+*/-*^ were comparable to those of wild-type (*Adamts19*^+*/*+^;*Mfrp*^+*/*+^), they were used as controls (Fig. S6). As expected, *Mfrp* mutant mice (*Adamts19*^+*/-*^;*Mfrp*^*-/-*^) exhibited reduced ocular axial length and VCD compared to *Adamts19*^+*/-*^; *Mfrp*^+*/-*^ control mice (Fig. 5C-E). Importantly, the ocular axial length and VCD of *Adamts19*^*-/-*^;*Mfrp*^*-/-*^ mice were significantly reduced compared to *Mfrp* mutant mice (*Adamts19*^+*/-*^;*Mfrp*^*-/-*^) (Fig. 5C-E). In addition, retinal thickness was increased in *Adamts19*^+*/-*^;*Mfrp*^*-/-*^ and *Adamts19*^*-/-*^;*Mfrp*^*-/-*^ mice compared to control *Adamts19*^+*/-*^; *Mfrp*^+*/-*^ mice (Fig. S7). These findings further support a role for the upregulation of retinal *Adamts19* expression being part of a compensatory mechanism triggered by impaired ocular axial growth.

**Figure 5.**
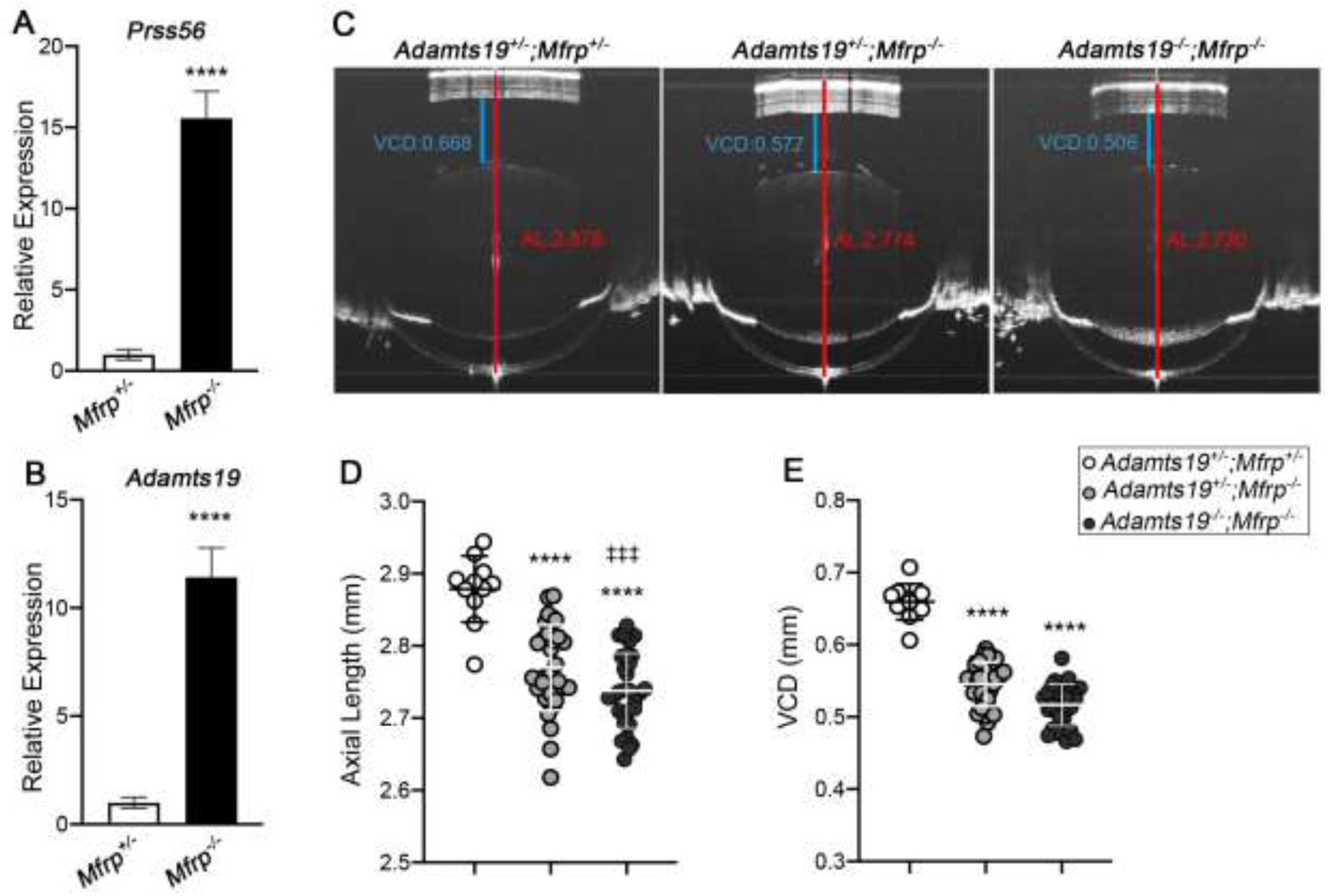
*Mfrp* mutant mice induced ocular size reduction is exacerbated by Adamts19 inactivation. (**A**-**B**) Histogram showing relative *Prss56 (***A**) and *Adamts19* (**B**) mRNA levels in *Mfrp*^+*/-*^ and *Mfrp*^*-/-*^ retina at P15. A significant increase in *Prss56 (***A**) and *Adamts19* (**B**) was detected in *Mfrp*^*-/-*^ compared to *Mfrp*^+*/-*^ retina, N>4/group. (**C**) Representative OCT images showing that ocular axial length (quantified in **D**) and VCD (quantified in **E**) are reduced in *Mfrp single mutant (Adamts19*^+*/-*^; *Mfrp*^*-/-*^) *and Adamts; Mfrp double mutant (Adamts1*^*-/-*^; *Mfrp*^*-/-*^) mice compared to control *(Adamts19*^+*/-*^; *Mfrp*^+*/-*^) eyes at P18, N>15 groups. Of note, the ocular axial length was significantly more reduced in *Adamts19*^*-/-*^;*Mfrp*^*-/-*^ than *Adamts19*^+*/-*^;*Mfrp*^*-/-*^ mice. Data are presented as mean ± SD. **\*\*\*\***p<0.0001 (compared to controls); ^‡‡‡^p<0.001 (compared to *Mfrp* single mutant), One way ANOVA.

### Inactivation of *Prss56 or Mfrp* prevents excessive ocular axial elongation in *Irbp* mutant mice

To further establish the role of PRSS56 and MFRP in ocular elongation, we tested the effects of *Prss56* and *Mfrp* inactivation in a mouse model of early-onset developmental myopia associated with excessive ocular axial growth caused by a null mutation in the gene coding for IRBP (Interphotoreceptor retinoid-binding protein)[24]. To this end, each of the *Prss56* and *Mfrp* mutant lines were crossed to *Irbp* mutant mice and biometric ocular assessment was conducted on their progeny. As expected, OCT analyses revealed that ocular axial length and VCD were significantly increased in *Irpb* single mutant mice (*Irbp*^*-/-*^;*Prss56*^+*/-*^ or *Irbp*^*-/-*^:*Mfrp*^+*/-*^) and significantly reduced in *Prss56 or Mfrp* single mutant mice (*Irbp*^+*/-*^;*Prss56*^*-/-*^ or *Irbp*^+*/-*^;*Mfrp*^*-/-*^,) compared to their respective controls (*Irbp*^+*/-*^;*Prss56*^+*/-*^ and *Irbp*^+*/-*^:*Mfrp*^+*/-*^ mice) (Fig. 6A-D). Inactivation of either *Prss56* or *Mfrp* prevented ocular axial elongation in *Irbp* mutant mice (*Irbp*^*-/-*^;*Prss56*^*-/-*^ and *Irbp*^*-/-*^;*Mfrp*^*-/-*^, respectively) (Fig. 6A-D). Notably, ocular axial length and VCD were significantly reduced in both double mutant lines (*Irbp*^*-/-*^;*Prss56*^*-/-*^ and *Irbp*^*-/-*^;*Mfrp*^*-/-*^) compared to their respective control littermates (*Irbp*^+*/-*^;*Prss56*^+*/-*^ and *Irbp*^+*/-*^;*Mfrp*^+*/-*^, respectively) and were comparable to those observed in *Prss56* and *Mfrp* single mutant mice (*Irbp*^+*/-*^;*Prss56*^*-/-*^ and *Irbp*^+*/-*^;*Mfrp*^*-/-*^, respectively) (Fig. 6A-D and Fig. S8 A, C). In addition, while retinal thickness was increased in both *Prss56* and *Mfrp* single mutant mice (*Irbp*^+*/-*^;*Prss56*^*-/-*^ and *Irbp*^+*/-*^;*Mfrp*^*-/-*^), it was significantly reduced in *Irpb* single mutant mice (*Irbp*^*-/-*^;*Prss56*^+*/-*^ or *Irbp*^*-/-*^;*Mfrp*^+*/-*^) compared to control littermates (*Irbp*^+*/-*^;*Prss56*^+*/-*^ and *Irbp*^+*/-*^:*Mfrp*^+*/-*^, respectively) (Fig. S8B, D). Interestingly, the retinal thickness of *Irbp*^*-/-*^; *Prss56*^*-/-*^ and *Irbp*^*-/-*^;*Mfrp*^*-/-*^ mice was comparable to that of their *Prss56* and *Mfrp* single mutant littermates (*Irbp*^+*/-*^;*Prss56*^*-/-*^ and *Irbp*^+*/-*^;*Mfrp*^*-/-*^, respectively) (Fig. S8B, D). Together, these findings demonstrate that the excessive ocular elongation observed in *Irbp*^*-/-*^ mice is dependent on PRSS56 and MFRP functions.

**Figure 6.**
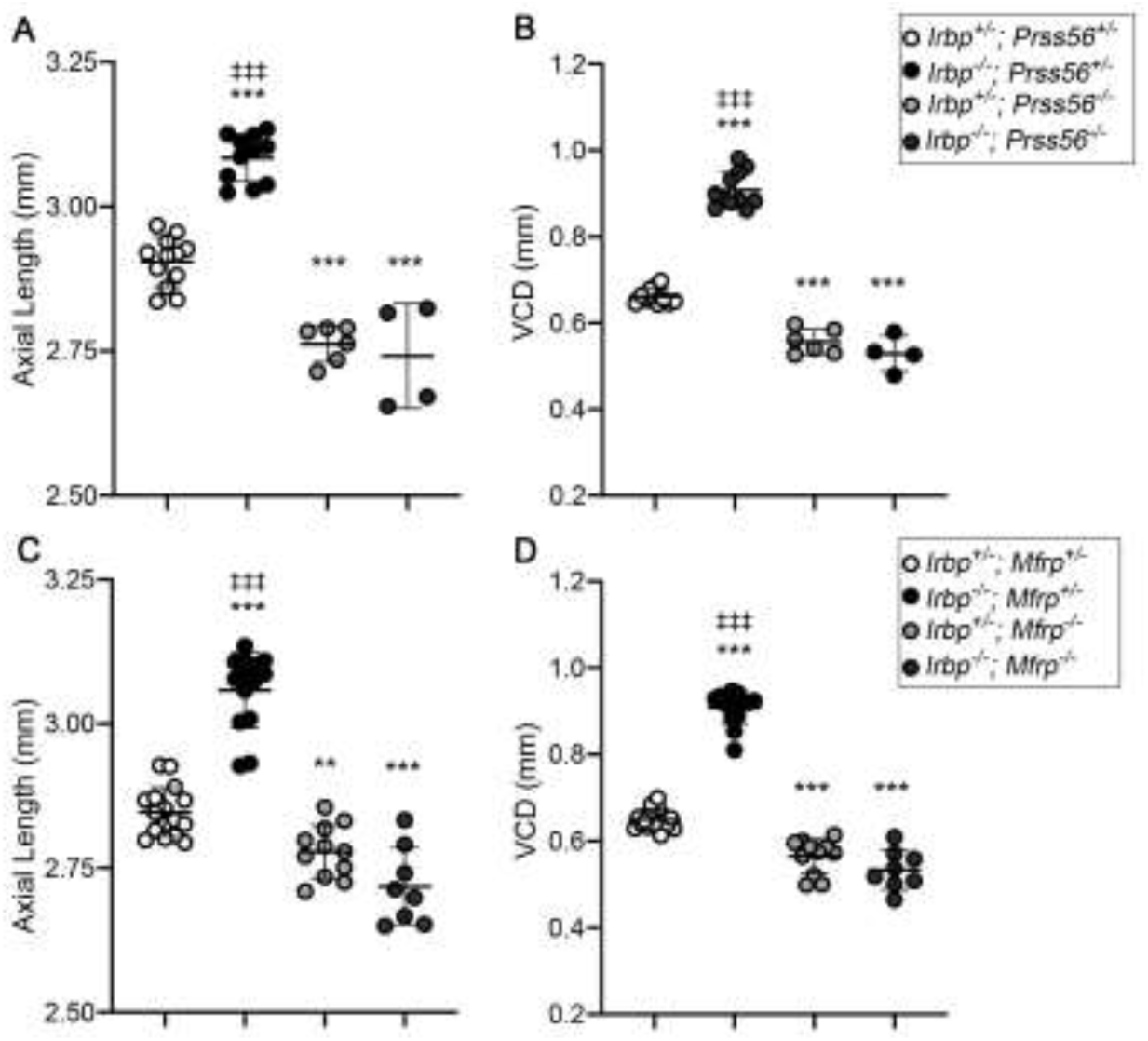
Ocular axial length elongation in *Irbp* mutant mice is dependent on PRSS56 or MFRP. (**A**-**D**) Scatter plot showing ocular axial length (**A** and **C**) and vitreous chamber depth (VCD) (**B** and **D**) values in *Irbp* mutant or control mice carrying a heterozygous or recessive mutation in *Prss56* or *Mfrp*: *Irbp* single mutants (*Irpb*^*-/-*^; *Prss56*^+*/-*^ in **A** and **B**, and *Irpb*^*-/-*^;*Mfrp*^+*/-*^ in **C** and **D**), *Irbp;Prss56* double mutant mice *(Irpb*^*-/-*^;*Prss56*^*-/-*^ in **A** and **B**), *Irpb; Mfrp* double mutant mice (*Irpb*^*-/-*^;*Mfrp*^*-/-*^ in **C** and **D**) and *Prss56* and *Mfrp* single mutant mice (*Irpb*^+*/-*^;*Prss56*^*-/-*^ in **A** and **C** and *Irpb*^+*/-*^; *Mfrp*^*-/-*^ in **B** and **D** respectively). Biometric analyses revealed that significant ocular axial elongation in *Irbp* single mutant mice (*Irpb*^*-/-*^;*Prss56*^+*/-*^ or *Irpb*^*-/-*^;*Mfrp*^+*/-*^) compared to control mice (*Irpb*^+*/-*^;*Prss56*^+*/-*^ or *Irpb*^+*/-*^;*Mfrp*^+*/-*^, respectively) that contrasts with the ocular axial length reduction observed in *Prss56* and *Mfrp* single mutant mice (*Irpb*^+*/-*^;*Prss56*^*-/-*^ and *Irpb*^+*/-*^; *Mfrp*^*-/-*^, respectively). Notably, *Prss56* and *Mfrp* inactivation prevented the ocular axial length elongation observed in *Irbp* mutant mice, as the ocular biometry of *Irpb*^*-/-*^;*Prss56*^*-/-*^ and *Irpb*^*-/-*^; *Mfrp*^*-/-*^ are comparable to *Prss56* or *Mfrp* single mutants, respectively (*Irpb*^+*/-*^;*Prss56*^*-/-*^ and *Irpb*^+*/-*^; *Mfrp*^*-/-*^, respectively). Overall, they suggest that PRSS56 and MFRP functions are required to induce ocular axial length elongation in *Irbp* mutant mice. Data are presented as mean ± SD (**A-D**) and as fold expression relative to wild-type (mean ± SEM), N=2 to 10/group (**E-F**). **\*\***p<0.01; **\*\*\***p<0.0001 (compared to controls); ^‡‡‡^p<0.001 (compared to double mutant *Irpb*^*-/-*^;*Prss56*^*-/-*^ and *Irpb*^*- /-*^; *Mfrp*^*-/-*^ mice), One way Anova.

## DISCUSSION

The molecular and cellular mechanisms involved in ocular axial growth and emmetropization are poorly understood. Previous studies have identified *PRSS56* and *MFRP* mutations as a major cause of nanophthalmos, a condition characterized by severe ocular size reduction and extreme hyperopia, suggesting that these factors play a critical role in ocular axial growth[3-6]. Consistent with this, *Prss56* and *Mfrp* mutant mice recapitulate the characteristic pathophysiological features of nanophthalmos, i.e. exhibit reduced ocular axial length and hyperopia[3, 13, 21]. Here, we use complementary genetic approaches in *Prss56* and *Mfrp* mutant mouse models as a first step to elucidate the molecular and cellular factors playing a role in the ocular size regulation. Notably, we identified ADAMTS19 as a novel factor involved in ocular size regulation and demonstrate that the upregulation of retinal *Adamts19* expression is part of a protective molecular response to impaired ocular growth. Also, we use a complementary strategy to show that inactivation of *Prss56* or *Mfrp* prevents excessive ocular elongation in a mouse model of early-onset developmental myopia caused by a null mutation in *Irpb*. Overall, our findings suggest that PRSS56 and MFRP are not only necessary for supporting ocular axial elongation under normal conditions, but also in the context of childhood-onset high myopia.

Gene expression profiling led us to the identification of PRSS56 and ADAMTS19 two secreted serine proteases, whose expression is altered in the retina of mouse model recapitulating features of nanophthalmos. We have previously reported that increased retinal expression of *Prss56* was a key molecular feature of *Prss56* mutant mice exhibiting a reduction in ocular axial length [13]. Here, we show that *Adamts19* expression is also upregulated in the retina of *Prss56* mutant mice. Importantly, taking advantage of the *Egr1;Prss56* double mutant mouse model in which *Egr1* inactivation rescues the ocular size reduction caused by loss of PRSS56 function, we demonstrate that the increased expression of retinal *Adamts19* results from ocular size reduction and is not a direct consequence of *Prss56* mutation per se. Further to support this finding, we show that retinal *Prss56* and *Adamts19* mRNA levels are also upregulated in an independent mouse model of nanophthalmos caused by a null *Mfrp* mutation. Importantly, we show that ADAMTS19 is not required for ocular axial growth during normal development, however, *Adamts19* inactivation exacerbates the reduction in ocular axial length and VCD in both *Prss56* and *Mfrp* mutant mouse models. Collectively, these findings indicate that the upregulation of retinal *Prss56* and *Adamst19* expression constitutes a protective response to overcome impaired ocular axial growth in two distinct mouse models of nanophthalmos. Since both PRSS56 and ADAMTS19 belong to the family of secreted serine-protease, it raises the possibility that they likely have overlapping or redundant function(s) and share the same substrate(s), which might explain the compensatory effect of ADAMTS19 on ocular elongation in mutant mice lacking PRSS56. Interestingly a recent study has found an association between genetic variant near *Adamts19* and ocular axial length, making our findings in mice relevant to human ocular size regulation [25].

Using a complementary genetic approach, we demonstrate that *Prss56* and *Mfrp* inactivation prevents the excessive ocular axial growth observed in a mouse model of early-onset high myopia caused by a null mutation in *Irbp [24]*. In the currently accepted model of ocular axial elongation, signals originating from the retina must first be relayed to the RPE before being transmitted to the choroid and subsequently to the sclera to induce scleral ECM remodeling and ocular axial growth [14, 18]. Importantly, the expression pattern of *Prss56* and *Adamts19* in the retina and that of *Mfrp* in the RPE are consistent with a central role for the retina and RPE in promoting ocular axial growth [13, 16]. More specifically, the cellular localization of *Prss56 and Adamts19* highlights the importance of Müller glia in mediating crosstalk between the retina and RPE. Interestingly, IRBP is localized in the interphotoreceptor matrix, a layer occupying the subretinal space juxtaposing the retinal photoreceptor cells and RPE. Thus, IRBP along with PRSS56 and MFRP may be part of a signaling network that not only connects the retina and RPE but also facilitates the flow of information, which are integral to ocular growth regulation. The increased expression of retinal *Prss56* in *Mfrp* mutant mice further lends support to the existence of potential crosstalk between Müller glia and RPE in the regulation of ocular axial growth. Overall, our findings suggest that PRSS56 and MFRP are critical for ocular axial growth during early developmental stages. Furthermore, as PRSS56 and MFRP are localized in the Müller glia and RPE respectively, they point towards a role for the interplay between Müller glia and RPE in ocular axial growth regulation.

We have shown previously that the ocular expression of *Prss56* is restricted to the retina, and predominantly observed in Müller glia [13]. Also, Prss56 upregulation was seen in retinal Müller glia of *Mfrp* mutant mice [23]. Here, we show that *Adamst19* expression is specifically detected in the INL of the *Prss56* mutant retina, a region where the cell body of Müller glia soma is found. These findings suggest that impaired ocular growth triggers the activation of a transcriptional program in retinal Müller glia leading to increased expression of *Prss56* and *Adamts19*, two genes encoding secreted serine proteases. Müller cells have been postulated to play a role in the detection of subtle changes in retinal structure due to mechanical stretching of their long processes or side branches[26, 27]. Reduction in ocular size may alter the structural and mechanical properties of the retina that are sensed by Müller glia triggering transcriptional activation of factors participating in the regulation of ocular axial growth [27].

Since genetic variants of PRSS56 and MFRP are also associated with common forms of myopia[11, 17], it raises the possibility of whether a nexus between Müller glia and RPE may have a broader role that contributes to vision-guided postnatal ocular growth. Also, given that loss of PRSS56 and MFRP function causes a reduction in ocular length[6, 13], it is plausible their noncoding variants associated with myopia may cause an increase in gene expression and thus, act via gain of function mechanism thereby contributing to an opposite phenotype characterized by an increase in ocular axial length. Furthermore, supporting the role of PRSS56 in myopia pathogenesis, a recent study in marmoset has shown an increase in retinal expression of Prss56 in response to minus lens-induced axial elongation/myopic compared to the control eyes[28]. Future efforts will focus on determining the cellular and molecular basis of the potential crosstalk between Müller glia and RPE in the regulation of ocular axial growth and their relevance to axial elongation in the context of myopia.

In summary, we identify ADAMTS19 as a novel factor involved in ocular size regulation and use a distinct mouse model of hyperopia/reduced ocular size and myopia/ excessive ocular growth to describe a regulatory genetic network playing a central role in regulating eye growth during development and disease. Collectively, these findings raise the possibility that modulation of *Adamts19* expression could be part of the general adaptive mechanism needed for regulating ocular axial growth. Furthermore, they suggest that PRSS56 and MFRP are indispensable for normal and aberrant ocular axial growth as a consequence of mutation in *Irbp* and point towards *Prss56* and *Mfrp* likely being part of a sequential pathway necessary for supporting ocular elongation.

## AUTHOR CONTRIBUTIONS

SK and KSN conceived and designed the study. SK, CL-D, SP, and YZ performed experiments. SK, CL-D, YZ, and KSN interpreted the results of analyses on mouse study.SK, CL-D, KSN, contributed to the drafting of the original manuscript. SK, CL-D, and KSN critically reviewed the manuscript.

## ACKNOWLEDGMENTS

This work was made possible, in part, by NEI P30 EY002162 - Core Grant for vision research (UCSF, Ophthalmology), Research to Prevent Blindness unrestricted grant (UCSF, Ophthalmology) and William and Mary Greve Special Scholar Award, That Man May See Inc, Research Evaluation and Allocation Committee (REAC)-Tidemann fund, and Marin Community Foundation-Kathlyn McPherson Masneri and Arno P. Masneri Fund (KSN) as well as by the Knight Templar Eye Foundation Career Starter Award (SK). The authors would like to acknowledge Ms. Vivian Chi and Mr. Yusef Seymens with mouse genotyping assay and general laboratory care.

## MATERIALS AND METHODS

### Animals

All experiments were conducted in compliance with protocols approved by the Institutional Animal Care and Use Committee at University of California San Francisco (IACUC) (Protocols # AN181358-01D) and following the guidelines from the Association for Research in Vision and Ophthalmology’s statement on the use of animals in ophthalmic research. Animals were given access to food and water ad libitum and housed under controlled conditions including 12-h light/dark cycle per the National Institutes of Health guidelines. Both male and female mice were used in all experiments and no differences were observed between sexes, all comparisons were made between littermates to minimalize variability.

### Mouse lines

1. ***Prss56***^***-/-***^ :(C57BL/6, Cg-*Prss56*^*glcr4*^/SjJ) – Mice carrying ENU induced mutation in *Prss56* causing truncation of PRSS56 protein at its C-terminal region [3].
2. ***Prss56***^***cre/cre***^ : (C57BL/6.Cg-Prss56tm(cre)) – Mice carrying a null allele of *Prss56* in which the exon1 of *Prss56* is replaced by CRE recombinase sequence[13, 29].
3. ***Egr1***^***-/-***^ :(C57BL/6. Egr1^tm1Jmi^/J) - Egr1 mutant mice: C57BL/6. Egr1tm1Jmi/J, the targeted mutation by insertion of a PGK-neo cassette introduces stop codon resulting in protein truncation upstream of the DNA-binding domain[30].
4. ***Adamts19***^***-/-***^ :(*Adamts19*^*tm4a(EUCOMM)Wtsi)*^) - A conditional *Adamts19* knockout mouse with *LoxP* sites flanking exon3. *E*xcision of the LoxP sites by ubiquitously expressed CRE recombinase driven by beta-actin promoter leads to the generation of a knockout allele of *Adamts19* (Supplementary Fig.2 A&B).
5. ***Mfrp***^***-/-***^ :(B6.C3Ga-Mfrp^rd6^/J): The mouse strain is homozygous for rd6 exhibiting retinal degeneration around four weeks during retinal developmental phase [23].
6. ***Irbp***^***-/-***^ :(B6.129P2-Rbp3^tm1Gil^/J): A knockout mouse model *Irbp(*Interstitial retinal binding protein 3) gene. This mouse line carries a targeted mutation for the *Rbp3* gene where the promoter and Exon 1 have been replaced by a NEO selection cassette rendering *Irbp* protein inactive.

PCR genotyping of all mouse strains was performed on genomic DNA obtained from tail biopsies digested with Proteinase K (Sigma, St. Louis, MO, USA) using primers listed in Table S1.

### Ocular Biometry

Ocular biometry was performed using Envisu R4300 spectral-domain optical coherence tomography (SD-OCT, Leica/Bioptigen Inc., Research Triangle Park, NC, USA). Measurements of various ocular parameters including axial length, vitreous chamber depth (VCD), anterior chamber depth (ACD), lens diameter and retinal thickness were performed on mice anesthetized with ketamine/xylazine (100 mg/kg and 5mg/kg, respectively; intraperitoneal (IP)) following pupil dilation as described previously[13].

### Quantitative polymerase chain reaction (qPCR)

For qPCR analysis of gene expression, eyes were enucleated and retinas were immediately dissected and total RNA was extracted from mouse retinal tissue using Qiagen RNeasy Mini Kit as per manufacturers protocol (Qiagen, Valencia, CA, USA) and reverse-transcribed using iScript cDNA Synthesis Kit (Bio-Rad, Hercules, CA, USA) and primer sets listed in Table S2. qPCR was performed on Bio-Rad C1000 Thermal Cycler/CF96 Real-Time System using SSOAdvanced™SYBR Green^®^ Supermix (Bio-Rad, Hercules, CA, USA). Briefly, 100ng of cDNA and 10uM primers were used per reaction in a final volume of 20ul. Each cycle consisted of denaturation at 95°C for 15s, followed by annealing at 60°C for 15s, extension 72°C for 30s for a total of 39 cycles. All the experiments were run as technical duplicates and a minimum of three biological replicates were used per group. The relative expression level of each gene was normalized to housekeeping genes (*Actinβ* and *Mapk1*) and analyzed using the CFX Maestro software (Bio-Rad, Hercules, CA, USA).

### *In situ* hybridization

Mice were transcardially perfused with ice-cold RNase-free PBS followed by 4% PFA (in RNase-free PBS). Eyes were enucleated post-fixed in RNAse-free 4% PFA, cryoprotected in 20% sucrose, and embedded in OCT and sectioned within 24 hours for *in situ* hybridization. QuantiGene View RNA (Affymetrix, Santa Clara, CA, USA). *In situ* hybridization was performed according to the manufacturer protocol. Briefly, 12µm cryosections were fixed overnight in 4% PFA, dehydrated through a graded series of ethanol, were subjected to 2X protease digestion for 10 minutes, fixed and hybridized with probe sets against *Adamts19* (NM_175506 (*Adamts19*), TYPE1, high sensitivity with 40∼50 bp DNAs) for 3 hours at 40°C using a ThermoBrite system (Abbott Molecular, Des Plaines, IL, USA). Cryosections were then washed and subject to signal amplification and detection using a fast red substrate, counterstained and mounted for subsequent imaging. Fluorescent images were acquired using an AxioImager M1 microscope equipped with an MRm digital camera and AxioVision software, with an LSM700 confocal microscope and Zen software (Carl Zeiss Microscopy, LLC, Germany).

### Statistical analyses

Statistical comparisons between mutant and control groups or between multiple experimental groups at a given age were performed using two-tailed unpaired Student’s t-test and one-way ANOVA, respectively, using Prism statistical software (version 6.02, GraphPad Software, San Diego, CA). A p-value of < (0.05, 0.01 and 0.001) was considered significant.

## REFERENCES

1. Carricondo PC, Andrade T, Prasov L, Ayres BM, Moroi SE. Nanophthalmos: A Review of the Clinical Spectrum and Genetics. J Ophthalmol. 2018;2018:2735465. doi: 10.1155/2018/2735465. PubMed PMID: 29862063; PubMed Central PMCID: PMCPMC5971257.

2. Siggs OM, Awadalla MS, Souzeau E, Staffieri SE, Kearns LS, Laurie K, et al. The genetic and clinical landscape of nanophthalmos and posterior microphthalmos in an Australian cohort. Clin Genet. 2020;97(5):764–9. doi: 10.1111/cge.13722. PubMed PMID: 32052405.

3. Nair KS, Hmani-Aifa M, Ali Z, Kearney AL, Ben Salem S, Macalinao DG, et al. Alteration of the serine protease PRSS56 causes angle-closure glaucoma in mice and posterior microphthalmia in humans and mice. Nature genetics. 2011;43(6):579-84. Epub 2011/05/03. doi: 10.1038/ng.813. PubMed PMID: 21532570.

4. Gal A, Rau I, El Matri L, Kreienkamp HJ, Fehr S, Baklouti K, et al. Autosomal-recessive posterior microphthalmos is caused by mutations in PRSS56, a gene encoding a trypsin-like serine protease. American journal of human genetics. 2011;88(3):382-90. Epub 2011/03/15. doi: 10.1016/j.ajhg.2011.02.006. PubMed PMID: 21397065; PubMed Central PMCID: PMC3059417.

5. Orr A, Dube MP, Zenteno JC, Jiang H, Asselin G, Evans SC, et al. Mutations in a novel serine protease PRSS56 in families with nanophthalmos. Molecular vision. 2011;17:1850-61. Epub 2011/08/19. PubMed PMID: 21850159; PubMed Central PMCID: PMC3137557.

6. Sundin OH, Leppert GS, Silva ED, Yang J-M, Dharmaraj S, Maumenee IH, et al. Extreme hyperopia is the result of null mutations in MFRP, which encodes a Frizzled-related protein. Proceedings of the National Academy of Sciences of the United States of America. 2005;102(27):9553-8. PubMed PMID: Medline:15976030.

7. Awadalla MS, Burdon KP, Souzeau E, Landers J, Hewitt AW, Sharma S, et al. Mutation in TMEM98 in a large white kindred with autosomal dominant nanophthalmos linked to 17p12-q12. JAMA Ophthalmol. 2014;132(8):970–7. doi: 10.1001/jamaophthalmol.2014.946. PubMed PMID: 24852644.

8. Cross SH, McKie L, Hurd TW, Riley S, Wills J, Barnard AR, et al. The nanophthalmos protein TMEM98 inhibits MYRF self-cleavage and is required for eye size specification. PLoS genetics. 2020;16(4):e1008583. doi: 10.1371/journal.pgen.1008583. PubMed PMID: 32236127; PubMed Central PMCID: PMCPMC7153906.

9. Garnai SJ, Brinkmeier ML, Emery B, Aleman TS, Pyle LC, Veleva-Rotse B, et al. Variants in myelin regulatory factor (MYRF) cause autosomal dominant and syndromic nanophthalmos in humans and retinal degeneration in mice. PLoS genetics. 2019;15(5):e1008130. doi: 10.1371/journal.pgen.1008130. PubMed PMID: 31048900; PubMed Central PMCID: PMCPMC6527243.

10. Almoallem B, Arno G, De Zaeytijd J, Verdin H, Balikova I, Casteels I, et al. The majority of autosomal recessive nanophthalmos and posterior microphthalmia can be attributed to biallelic sequence and structural variants in MFRP and PRSS56. Sci Rep. 2020;10(1):1289. doi: 10.1038/s41598-019-57338-2. PubMed PMID: 31992737; PubMed Central PMCID: PMCPMC6987234.

11. Hysi PG, Choquet H, Khawaja AP, Wojciechowski R, Tedja MS, Yin J, et al. Meta-analysis of 542,934 subjects of European ancestry identifies new genes and mechanisms predisposing to refractive error and myopia. Nature genetics. 2020;52(4):401–7. doi: 10.1038/s41588-020-0599-0. PubMed PMID: 32231278; PubMed Central PMCID: PMCPMC7145443.

12. Siegwart JT, Jr., Norton TT. Perspective: how might emmetropization and genetic factors produce myopia in normal eyes? Optometry and vision science : official publication of the American Academy of Optometry. 2011;88(3):E365-72. PubMed PMID: Medline:21258261.

13. Paylakhi S, Labelle-Dumais C, Tolman NG, Sellarole MA, Seymens Y, Saunders J, et al. Muller glia-derived PRSS56 is required to sustain ocular axial growth and prevent refractive error. PLoS genetics. 2018;14(3):e1007244. doi: 10.1371/journal.pgen.1007244. PubMed PMID: 29529029; PubMed Central PMCID: PMCPMC5864079.

14. Stone RA, Pardue MT, Iuvone PM, Khurana TS. Pharmacology of myopia and potential role for intrinsic retinal circadian rhythms. Experimental eye research. 2013;114:35-47. Epub 2013/01/15. doi: 10.1016/j.exer.2013.01.001. PubMed PMID: 23313151; PubMed Central PMCID: PMC3636148.

15. Morgan IG, Ohno-Matsui K, Saw SM. Myopia. Lancet. 2012;379(9827):1739-48. Epub 2012/05/09. doi: 10.1016/S0140-6736(12)60272-4. PubMed PMID: 22559900.

16. Sundin OH, Dharmaraj S, Bhutto IA, Hasegawa T, McLeod DS, Merges CA, et al. Developmental basis of nanophthalmos: MFRP Is required for both prenatal ocular growth and postnatal emmetropization. Ophthalmic Genet. 2008;29(1):1-9. PubMed PMID: Medline:18363166.

17. Tedja MS, Wojciechowski R, Hysi PG, Eriksson N, Furlotte NA, Verhoeven VJM, et al. Genome-wide association meta-analysis highlights light-induced signaling as a driver for refractive error. Nature genetics. 2018;50(6):834-+. doi: 10.1038/s41588-018-0127-7. PubMed PMID: WOS:000433621000011.

18. Pardue MT, Stone RA, Iuvone PM. Investigating mechanisms of myopia in mice. Experimental eye research. 2013;114:96-105. Epub 2013/01/12. doi: 10.1016/j.exer.2012.12.014. PubMed PMID: 23305908; PubMed Central PMCID: PMC3898884.

19. Collery RF, Volberding PJ, Bostrom JR, Link BA, Besharse JC. Loss of Zebrafish Mfrp Causes Nanophthalmia, Hyperopia, and Accumulation of Subretinal Macrophages. Investigative ophthalmology & visual science. 2016;57(15):6805-14. Epub 2016/12/22. doi: 10.1167/iovs.16-19593. PubMed PMID: 28002843; PubMed Central PMCID: PMC5215506.

20. Fogerty J, Besharse JC. 174delG mutation in mouse MFRP causes photoreceptor degeneration and RPE atrophy. Investigative ophthalmology & visual science. 2011;52(10):7256-66. PubMed PMID: Medline:21810984.

21. Velez G, Tsang SH, Tsai YT, Hsu CW, Gore A, Abdelhakim AH, et al. Gene Therapy Restores Mfrp and Corrects Axial Eye Length. Sci Rep. 2017;7(1):16151. doi: 10.1038/s41598-017-16275-8. PubMed PMID: 29170418; PubMed Central PMCID: PMCPMC5701072.

22. Schippert R, Burkhardt E, Feldkaemper M, Schaeffel F. Relative axial myopia in Egr-1 (ZENK) knockout mice. Investigative ophthalmology & visual science. 2007;48(1):11–7. doi: 10.1167/iovs.06-0851. PubMed PMID: 17197510.

23. Soundararajan R, Won J, Stearns TM, Charette JR, Hicks WL, Collin GB, et al. Gene profiling of postnatal Mfrprd6 mutant eyes reveals differential accumulation of Prss56, visual cycle and phototransduction mRNAs. PloS one. 2014;9(10):e110299. Epub 2014/10/31. doi: 10.1371/journal.pone.0110299. PubMed PMID: 25357075; PubMed Central PMCID: PMC4214712.

24. Wisard J, Faulkner A, Chrenek MA, Waxweiler T, Waxweiler W, Donmoyer C, et al. Exaggerated eye growth in IRBP-deficient mice in early development. Investigative ophthalmology & visual science. 2011;52(8):5804-11. PubMed PMID: Medline:21642628.

25. Fan Q, Pozarickij A, Tan NYQ, Guo X, Verhoeven VJM, Vitart V, et al. Genome-wide association meta-analysis of corneal curvature identifies novel loci and shared genetic influences across axial length and refractive error. Commun Biol. 2020;3(1):133. doi: 10.1038/s42003-020-0802-y. PubMed PMID: 32193507; PubMed Central PMCID: PMCPMC7081241.

26. Bringmann A, Pannicke T, Grosche J, Francke M, Wiedemann P, Skatchkov SN, et al. Muller cells in the healthy and diseased retina. Progress in retinal and eye research. 2006;25(4):397-424. Epub 2006/07/15. doi: 10.1016/j.preteyeres.2006.05.003. PubMed PMID: 16839797.

27. Lindqvist N, Liu Q, Zajadacz J, Franze K, Reichenbach A. Retinal glial (Muller) cells: sensing and responding to tissue stretch. Investigative ophthalmology & visual science. 2010;51(3):1683-90. PubMed PMID: Medline:19892866.

28. Tkatchenko TV, Troilo D, Benavente-Perez A, Tkatchenko AV. Gene expression in response to optical defocus of opposite signs reveals bidirectional mechanism of visually guided eye growth. PLoS Biol. 2018;16(10):e2006021. doi: 10.1371/journal.pbio.2006021. PubMed PMID: 30300342; PubMed Central PMCID: PMCPMC6177118.

29. Jourdon A, Gresset A, Spassky N, Charnay P, Topilko P, Santos R. Prss56, a novel marker of adult neurogenesis in the mouse brain. Brain structure & function. 2016;221(9):4411-27. Epub 2015/12/25. doi: 10.1007/s00429-015-1171-z. PubMed PMID: 26701169.

30. Lee SL, Tourtellotte LC, Wesselschmidt RL, Milbrandt J. Growth and differentiation proceeds normally in cells deficient in the immediate early gene NGFI-A. J Biol Chem. 1995;270(17):9971-7. PubMed PMID: 7730380.

