## Supplement Figures and Table for "Identification of ADAMTS19 as a novel retinal factor involved in ocular growth regulation"

**SUPPLEMENTAL FIGURES**

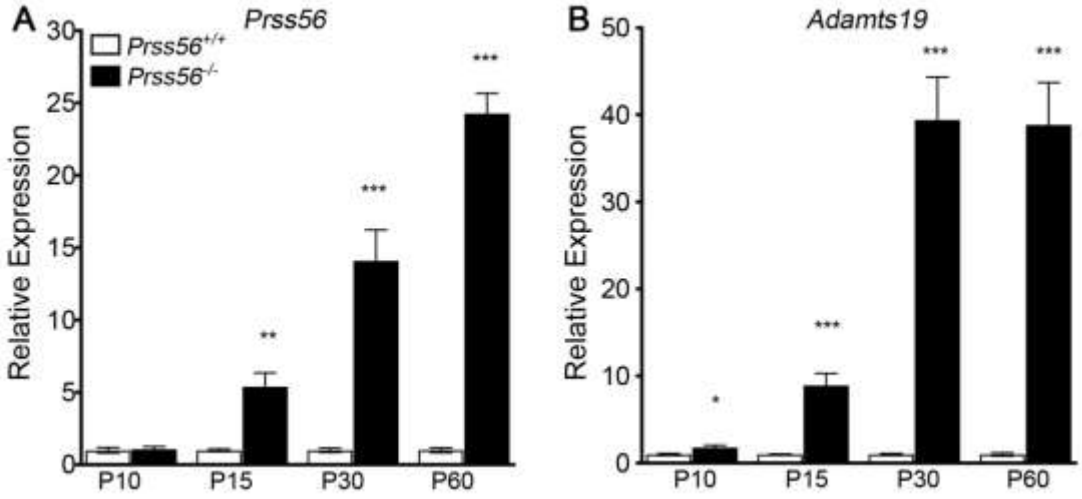


**Figure S1. *Prss56* and *Adamts19* expression in *Prss56*$^{-/-}$ retina across ages.** (**A**-**B**) Graphs showing quantification of *Prss56 (***A**) and *Adamts19* (**B**) mRNA levels using qPCR in wild-type and mutant retina at different developmental stages. A significant increase in *Adamts19* mRNA levels was detected as early as P10 in *Prss56* mutant retina (**B**), while upregulation in *Prss56* mRNA was first observed at P15 in the mutant retina (**A**). The magnitude of the increase of both *Prss56* and *Adamts19* expression became more pronounced with age in the mutant retina. *Prss56* and *Adamts19* expression were normalized to the expression of three housekeeping genes (*Hprt1*, *Actb1,* and *Mapk1).* Data are presented as fold expression relative to wild-type (mean ± SEM), N=4 to 6/group. *****p<0.05; ******p<0.01; *******p<0.001, t-test. The *Prss56* qPCR data in **A** were previously published in Figure 4 of Paylakhi et al. 2018 [13].

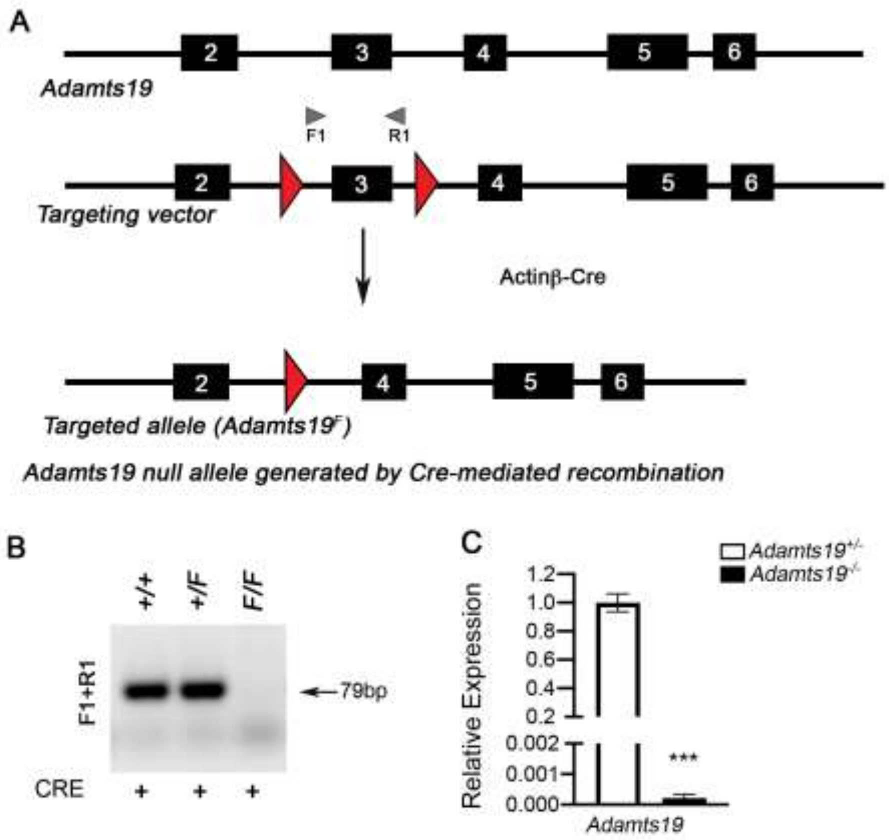


**Figure S2. Generation of *Adamts19* mutant mice allele.**

(**A**) The LoxP site flanks the exon 3 of the $Adamts19^{F}$ allele. In presence of Cre recombinase, the Adamts19 exon is deleted resulting in a frameshift mutation and premature stop codon, rendering the *Adamts19* catalytically inactive. (**B-C**) *Adamts19* exon 3 excisions was confirmed by PCR as well as qPCR. (**B)** PCR amplification of DNA from wild-type ($Adamts19^{+/+}$, lane 1), heterozygous ($Adamts19^{F/+,}$ lane 2), or homozygous ($Adamts19^{F/F}$) mice. PCR reactions were performed using primers amplifying regions of intron 2 and exon 3. The deletion of exon 3 from the $Adamts19^{F/F}$ allele gives no PCR product. using primers amplifying a region of intron 2 and exon 3 gives no PCR product from DNA from $Adamts19^{F/F}$ mice. (**C)** qPCR analysis employing primers amplifying exon 3 of A*damts19* further confirm that Adamts19 transcript from $Adamts19^{-/-}$;$Prss56^{-/-}$ retina lacks exon 3.

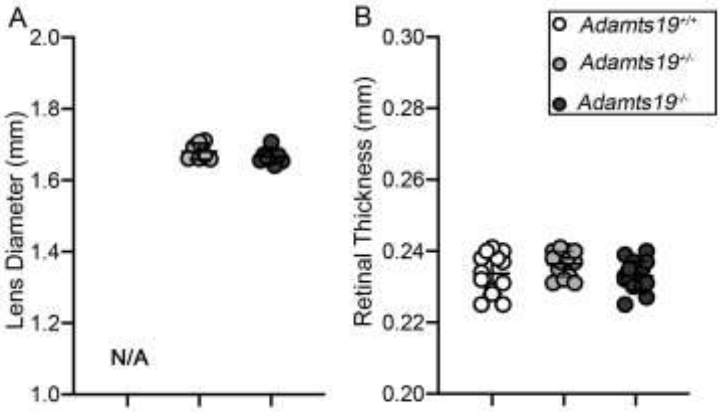


**Figure S3. Ocular biometric analysis in *Adamts19* mutant mice**

Histograms showing that ocular the lens diameter (**A**) and retinal thickness (**B**) were indistinguishable in *Adamts19*$^{-/-}$;*Prss56*$^{-/-}$, *Adamts19*$^{+/-}$;*Prss56*$^{-/-}$ and control *Adamts19*$^{+/-}$; *Prss56*$^{+/-}$ mice. Data are presented as mean ± SD, N$\geq$7/group. *p<0.05, One-way ANOVA.

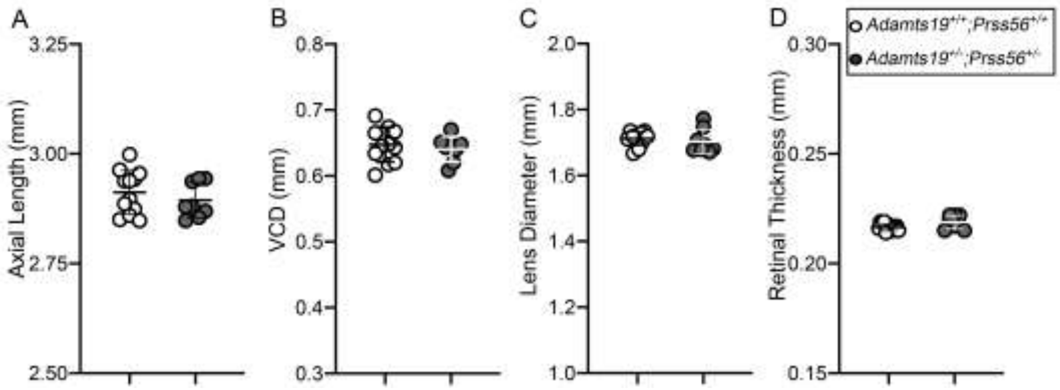


**Figure S4. Ocular biometric parameters are indistinguishable between wild-type and *Adamts19*$^{+/-}$*:Prss56*$^{+/-}$ mice.** (**A-D**) Histograms showing that all the ocular biometric parameters examined including axial length (**A**), vitreous (VCD) (**B**), lens diameters (**C**) and retinal thickness (**D**) are indistinguishable between wild-type (*Adamts19*$^{+/+}$*;Prss56*$^{+/+}$) and *Adamts19*$^{+/-}$*;Prss56*$^{+/-}$ mice at P18  Data are presented as mean ± SD, N$\geq$7/group. *p<0.05, One-way ANOVA.

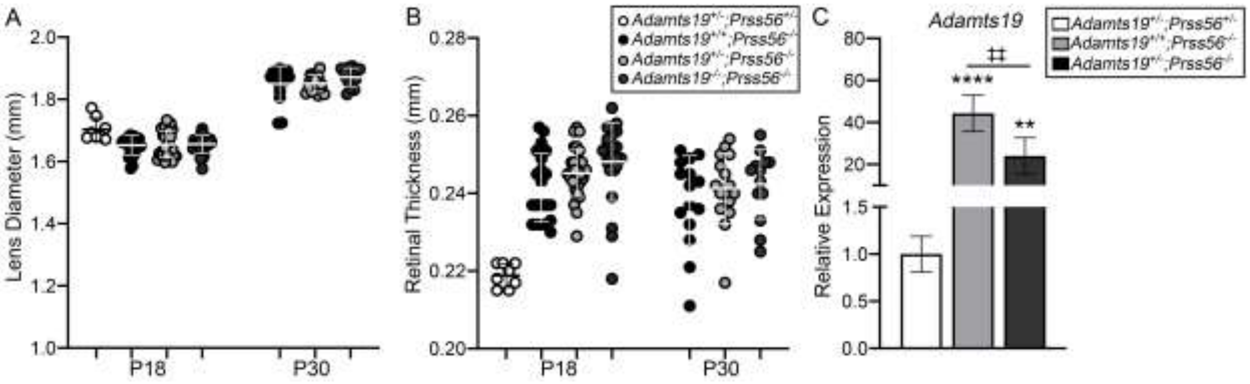


**S5. Ocular biometric analysis of *Adamts19$^{-/-}$;Prss56$^{-/-}$* mice.** (**A-B**) Histograms showing that retinal thickness (**A**) is modestly increased in *Adamts19$^{-/-}$;Prss56$^{-/-}$* mice compared to control *(Adamts19$^{+/-}$*; *Prss56$^{+/-}$*) mice at P18, but not at P30, while the lens diameter (**B**) is indistinguishable in *Adamts19$^{-/-}$;Prss56$^{-/-}$*, *Adamts19$^{+/-}$;Prss56$^{-/-}$* and control *(Adamts19$^{+/-}$*; *Prss56$^{+/-}$*) mice at both ages examined. (**C**) Histogram showing relative *Adamts19* mRNA levels AGE N>4/group. Data are presented as mean ± SD, N$\geq$13/group. *p<0.05, One-way ANOVA.

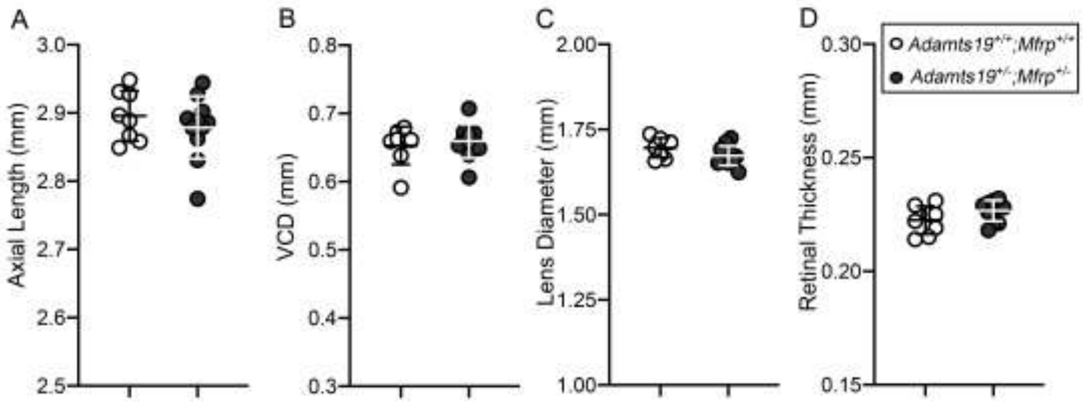


**Figure S6. Ocular biometric parameters are indistinguishable between wild-type and *Adamts19*$^{+/-}$*:Mfrp*$^{+/-}$ mice.** (**A-D**) Histograms showing that all the ocular biometric parameters examined including axial length (**A**), vitreous (VCD) (**B**), lens diameters (**C**), and retinal thickness (**D**) are indistinguishable between wild-type (*Adamts19*$^{+/+}$*;Mfrp*$^{+/+}$) and *Adamts19*$^{+/-}$*;Mfrp*$^{+/-}$ mice at P18. Data are presented as mean ± SD, N≥7/group. *p<0.05, One-way ANOVA.

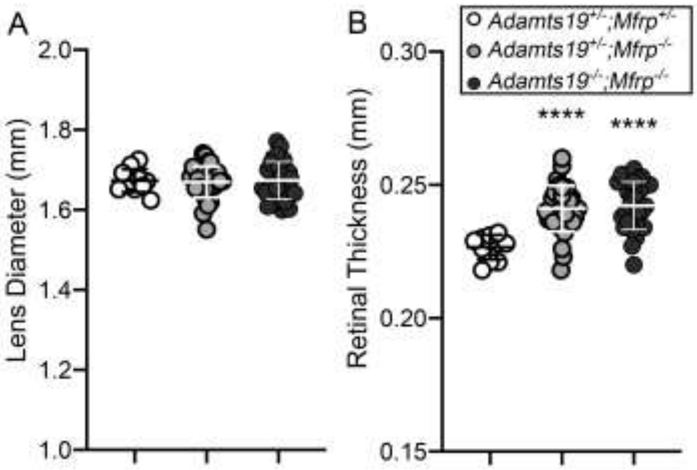


**Figure S7. Ocular biometric analysis of *Adamts19$^{-/-}$;Mfrp$^{-/-}$* mice.** (**A-C**) Histograms showing that the lens diameter (**A**) are indistinguishable in *Adamts19$^{+/-}$*; *Mfrp$^{-/-}$*, *Adamts1$^{-/-}$*; *Mfrp$^{-/-}$*, and control *(Adamts19$^{+/-}$*; *Mfrp$^{+/-}$*) mice, while the retinal thickness (**B**) is increased in both *Adamts19$^{+/-}$*; *Mfrp$^{-/-}$* and *Adamts1$^{-/-}$*;*Mfrp$^{-/-}$* mice compared to control *Adamts19$^{+/-}$*; *Mfrp$^{+/-}$* mice at P18. Data are presented as mean ± SD, N≥11/group. ****p<0.0001, One-way ANOVA.

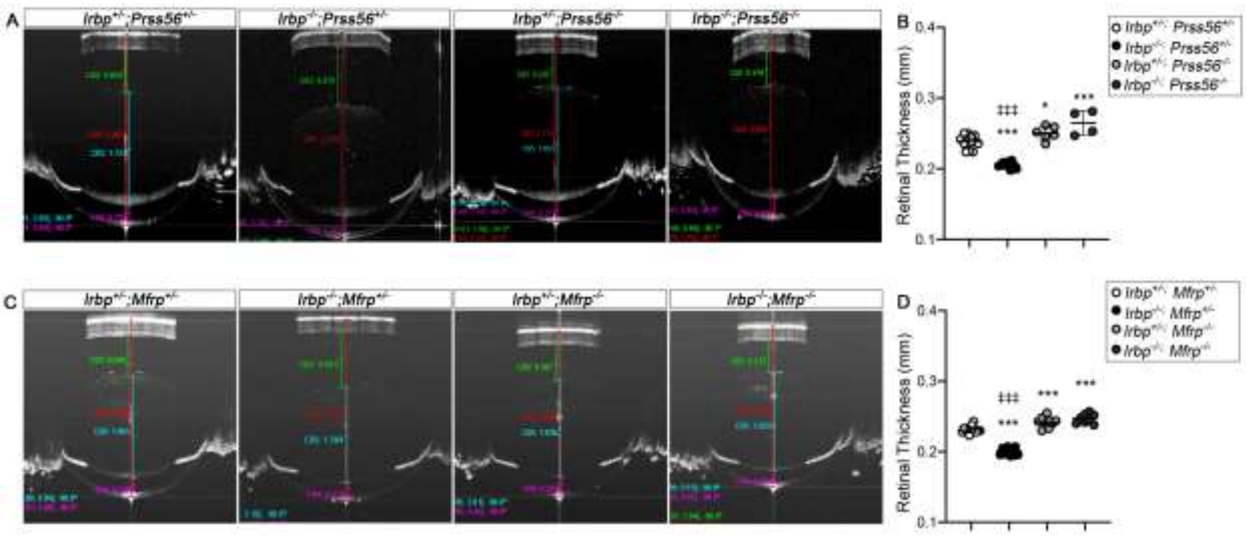


**Figure S8.**

**Ocular biometric analysis of *Irbp* mutant mice following *Prss56* inactivation**

**(A)** Representative OCT images showing that ocular axial length (quantified in Fig. 6A) and VCD (quantified in Fig. 6 B) are increased in *Irpb* mutant mice (*Irpb*$^{-/-}$*; Prss56*$^{+/-}$) and reduced in *Prss56* mutant mice compared to control *Irpb*$^{+/-}$*; Prss56*$^{+/-}$ control mice at P18. In contrast, retinal thickness (**B**) were reduced in *Irpb* mutant mice (*Irpb*$^{-/-}$*; Prss56*$^{+/-}$) and retinal thickness was increased in *Prss56* mutant and *Irbp;Prss56* double mutant mice (*Irpb*$^{+/-}$*; Prss56*$^{-/-}$ and *Irbp*$^{-/-}$*; Prss56*$^{-/-}$, respectively) compared to control *Irpb*$^{+/-}$*; Prss56*$^{+/-}$ mice. Data are presented as mean ± SD, N$\geq$4/group. ******p<0.01; *******p<0.0001 (compared to controls); ‡‡‡p<0.001 (compared to double mutant *Irpb*$^{-/-}$;*Prss56*$^{-/-}$ mice), One-way Anova.

**Biometric analysis of *Irbp* mutant mice following *Mfrp* inactivation** (**C**) Representative OCT images showing that ocular axial length (quantified in Fig. 6 C) and vitreous chamber depth (VCD, quantified in Fig. 6 D) are increased in *Irpb* mutant mice (*Irpb*$^{-/-}$*; Mfrp*$^{+/-}$) and reduced in *Mfrp* mutant and *Irbp;Mfrp* double mutant mice (*Irpb*$^{+/-}$*; Mfrp*$^{-/-}$ and *Irbp*$^{-/-}$*; Mfrp*$^{-/-}$, respectively) compared to control *Irpb*$^{+/-}$*; Mfrp*$^{+/-}$ mice at P18. In contrast, and retinal thickness (**D**) were reduced in *Irpb* mutant mice (*Irpb*$^{-/-}$*; Mfrp*$^{+/-}$) and retinal thickness was increased in *Mfrp* mutant and *Irbp;Mfrp* double mutant mice (*Irpb*$^{+/-}$*; Mfrp*$^{-/-}$ and *Irbp*$^{-/-}$*; Mfrp*$^{-/-}$, respectively) compared to control *Irpb*$^{+/-}$*; Mfrp*$^{+/-}$ mice. Data are presented as mean ± SD, N$\geq$8/group. ******p<0.01; *******p<0.0001 (compared to controls); ‡‡‡p<0.001 (compared to double mutant *Irpb*$^{-/-}$; *Mfrp*$^{-/-}$ mice), One-way Anova.

### Table S1. List of genotyping primers

| Allele (mouse strain) | Primer name | Primer sequence |
|---|---|---|
| ***Prss56***$^{glcr4}$ **(C57BL/6.Cg-Prss56 glcr4/SjJ)** | Prss56 glcr4 F1 | 5' TGGCTCCAGAAACCAAAGCCGGAA GAGCGCCCGGAAACAAAGAGT 3' |
| | Prss56 glcr4 F2 | 5' GCGGCGCCCGGAAACAAAAGGA 3' |
| | Prss56 glcr4 R | 5' TCCTGGAAGAGAGGGAGTGA 3' |
| ***Prss56***$^{Cre}$ **(C57Bl/6.Cg-Prss56tm)** | Prss56 Cre F | 5’ CAG GGC ATC GTT TCC CTG AG 3’ |
| | Prss56 Cre R WT | 5’ GAC AGG CGC GTG TAC AGT GG 3’ |
| | Prss56 Cre R Cre | 5’ CCA TGA GTG AAC GAA CCT GG 3’ |
| ***Egr1***$^{-/-}$ **(C57BL/6. Egr1tm1Jmi/J )** | EGR1 common R | 5' GGG CAC AGG GGA TGG GAA3 |
| | EGR1 MUT F | 5' AAC CGG CCC AGC AAG ACA 3' |
| | EGR1 WT F | 5' CTC GTG CTT TAC GGT ATC G 3' |
| ***Adamts19***$^{-/-}$ **(Adamts19tm4a(EUCOMM)Wtsi)** | Adamts19_244_F | 5’ AGA AGG GAA CAA ACA CAA CAA GTG 3’ |
| | Adamts19_244_R | 5’ AGT TAG CCT GAG CCT GTG TGG 3’ |
| | CAS_R1_Term | 5’ TCG TGG TAT CGT TAT GCG GCC 3’ |
| ***Mfrp***$^{-/-}$ **( B6.C3Ga-Mfrprd6/J )** | Mfrp F | 5’ CAC TAC CAC CCC AGC AAG GAC 3’ |
| | Mfrp R | 5’ CTT CTC CAG AGA GTG CCC TTG 3’ |
| ***Irbp***$^{-/-}$ **(B6.129P2-Rbp3tm1Gil/J)** | Common IRBP | 5’ CAT ATC CAC ACC TGC CAA CA 3’ |
| | MF IRBP | 5’ GCT ACT TCC ATT TGT CAC GTC C 3’ |
| | WT IRBP | 5’ GGA CCC ACA CCT GAA GAC AG3’ |
| ***Ubc-Cre*** **(C57Bl/6.Cg-Tg(UBC-Cre/ERT2)1Ejb)** | Cr1 | 5’ TGA TGA GGT TCG CAA GAA CC 3’ |
| | Cr2 | 5’ CCA TGA GTG AAC GAA CCT GG 3’ |

### Table S2. List of qPCR primers

| Gene | Forward Primer | Reverse Primer |
|---|---|---|
| ***House Keeping Genes*** | | |
| *Actb* | 5' CCCTGAGGAGCACCCTGTGC 3' | 5' GGCTGGGGTGTTGAAGGTCT 3' |
| *Hprt1* | 5' TGCCGAGGATTTGGAAAAAGTGT 3' | 5' GTGATGGCCTCCCATCTCCT 3' |
| *Mapk1* | 5' TTGAACAGGCTCTGGCCCAC 3' | 5' TGAATGGCGCTTCAGCAATGG 3' |
| ***Genes of Interest*** | | |
| *Prss56* | 5' ACCTGGACGCCCTAGACCTC 3' | 5' TGTTGGCAACGCCTTGATGT 3' |
| *Adamts19* | 5' `TGCTGAAGACAACGGCCTGA 3' | 5' CAGCACAGGATGGGTGGTCA 3' |
| ***Adamts19 Ex3*** | 5' GAGGACTTCCTATTCATTGAGCCA 3' | 5' CCTGTATAAACGGTGCGGGT 3' |
